# Decoding rRNA sequences for improved metagenomics of sylvatic mosquito species

**DOI:** 10.1101/2022.02.01.478639

**Authors:** Cassandra Koh, Lionel Frangeul, Hervé Blanc, Carine Ngoagouni, Sébastien Boyer, Philippe Dussart, Nina Grau, Romain Girod, Jean-Bernard Duchemin, Maria-Carla Saleh

**Author notes:** To whom correspondence should be addressed. Corresponding author: Maria-Carla Saleh. Philippe Dussart, Institut Pasteur de Madagascar, Antananarivo 101, Madagascar; Nina Grau, Sciences Economiques et Sociales de la Santé et Traitement de l’Information Médicale, Faculté de Médecine, Marseille 13005, France.

## Abstract

As mosquito-borne virus epidemics are often preceded by undetected spillover events, surveillance and virus discovery studies in non-urban mosquitoes informs pre-emptive and responsive public health measures. RNA-seq metagenomics is a popular methodology but it is constrained by overabundant rRNA. The lack of reference sequences for most mosquito species is a major impediment against physical and computational removal of rRNA reads.

We describe a strategy to assemble novel rRNA sequences from mosquito specimens, producing an unprecedented dataset of 234 full-length 28S and 18S rRNA sequences of 33 medically important species from countries with known histories of mosquito-borne virus circulation (Cambodia, the Central African Republic, Madagascar, and French Guiana). We also evaluate the utility of rRNA sequences as molecular barcodes relative to the mitochondrial *cytochrome* c *oxidase I* (COI) gene. We show that rRNA sequences can be used for species identification when COI sequences are ambiguous or unavailable, revealing evolutionary relationships concordant with contemporary mosquito systematics.

This expansion of the rRNA reference library improves mosquito RNA-seq metagenomics by permitting the optimization of species-specific rRNA depletion protocols for a broader species range and streamlined species identification by rRNA barcoding. In addition, rRNA barcodes could serve as an additional tool for mosquito taxonomy and phylogeny.

## INTRODUCTION

Mosquitoes top the list of vectors for arthropod-borne diseases, being implicated in the transmission of many human pathogens responsible for arboviral diseases, malaria, and lymphatic filariasis (WHO, 2017). Mosquito-borne viruses circulate in sylvatic or urban transmission cycles driven by different mosquito species with their own distinct host preferences. Although urban mosquito species are chiefly responsible for amplifying epidemics in dense human populations, non-urban (sylvatic and peri-urban) mosquitoes maintain the transmission of these viruses among forest-dwelling animal reservoir hosts and are implicated in spillover events when humans enter their ecological niches (Valentine et al., 2019). Given that mosquito-borne virus emergence is preceded by such spillover events, continuous surveillance and virus discovery in non-urban mosquitoes is integral to designing effective public health measures to pre-empt or respond to mosquito-borne viral epidemics.

Metagenomics on field specimens is the most powerful method in the toolkit for understanding mosquito-borne disease ecology through the One Health lens (Webster et al., 2016). With next-generation sequencing becoming more accessible, such studies have provided unprecedented insights into the interfaces among mosquitoes, their environment, and their animal and human hosts. However, working with lesser studied mosquito species poses several problems.

First, metagenomics studies based on RNA-seq are bedevilled by overabundant ribosomal RNAs (rRNAs). These non-coding RNA molecules comprise at least 80% of the total cellular RNA population (Gale & Crampton, 1989). Due to their length and their abundance, they are a sink for precious next generation sequencing reads, decreasing the sensitivity of pathogen detection unless depleted during library preparation. Yet the most common rRNA depletion protocols require prior knowledge of rRNA sequences of the species of interest as they involve hybridizing antisense oligos to the rRNA molecules prior to removal by ribonucleases (Fauver et al., 2019; Phelps et al., 2021) or by bead capture (Kukutla et al., 2013). Presently, reference sequences for rRNAs are limited to only a handful of species from three genera: *Aedes, Culex*, and *Anopheles* (Ruzzante et al., 2019). The lack of reliable rRNA depletion methods could deter mosquito metagenomics studies from expanding their sampling diversity, resulting in a gap in our knowledge of mosquito vector ecology. The inclusion of lesser studied yet medically relevant non-urban species is therefore imperative.

Second, species identification based on morphology is notoriously complicated between members of species subgroups. This is especially the case among *Culex* subgroups. Sister species are often sympatric and show at least some competence for a number of viruses, such as Japanese encephalitis virus, St Louis encephalitic virus, and Usutu virus (Nchoutpouen et al., 2019). Although they share many morphological traits, each of these species have distinct ecologies and host preferences, thus the challenge of correctly identifying vector species can affect epidemiological risk estimation for these diseases (Farajollahi et al., 2011). DNA molecular markers are often employed to a limited degree of success to distinguish between sister species (Batovska et al., 2017; Zittra et al., 2016).

To address the lack of full-length rRNA sequences in public databases, we sought to determine the 28S and 18S rRNA sequences of a diverse set of Old and New World non-urban (sylvatic and peri-urban) mosquito species from four countries representing three continents: Cambodia, the Central African Republic, Madagascar, and French Guiana. These countries, due to their proximity to the equator, contain high mosquito biodiversity (Foley et al., 2007) and have had long histories of mosquito-borne virus circulation. Increased and continued surveillance of local mosquito species could lead to valuable insights on mosquito virus biogeography. Using a unique score-based read filtration strategy to remove interfering non-mosquito rRNA reads for accurate *de novo* assembly, we produced a dataset of 234 novel full-length 28S and 18S rRNA sequences from 33 mosquito species, 30 of which have never been recorded before.

We also explored the functionality of 28S and 18S rRNA sequences as molecular barcodes by comparing their performance to that of the mitochondrial *cytochrome* c *oxidase subunit I* (COI) gene for molecular taxonomic and phylogenetic investigations. The COI gene is the most widely used DNA marker for molecular species identification and forms the basis of the Barcode of Life Data System (BOLD) (Hebert et al., 2003; Ratnasingham & Hebert, 2007). Presently, full-length rRNA sequences are much less represented compared to other molecular markers. We hope that our sequence dataset, with its species diversity and eco-geographical breadth, and the assembly strategy we describe would further facilitate the use of rRNA as barcodes. In addition, this dataset enables the design of species-specific oligos for cost-effective rRNA depletion for a broader range of mosquito species and streamlined molecular species diagnostics during RNA-seq.

## RESULTS

### Poor rRNA depletion using a non-specific depletion method

During library preparations of mosquito samples for RNA-seq, routinely used methods for depleting rRNA are commercial kits optimised for human or mice samples (Belda et al., 2019; Bishop-Lilly et al., 2010; Chandler et al., 2015; N. Kumar et al., 2012; Weedall et al., 2015; Zakrzewski et al., 2018) or through 80–100 base pair antisense probe hybridisation followed by ribonuclease digestion (Fauver et al., 2019; Phelps et al., 2021). In cases where the complete reference rRNA sequence of the target species is not known, oligos would be designed based on the rRNA sequence of the closest related species (25, this study). These methods should deplete the conserved regions of rRNA sequences. However, the variable regions remain at abundances high enough to compromise RNA-seq output. In our hands, we have found that using probes designed for the *Ae. aegypti* rRNA sequence followed by RNase H digestion according to the protocol published by Morlan *et al*. (2012) produced poor depletion in *Ae. albopictus*, and in Culicine and Anopheline species (Figure 1A). Additionally, the lack of reference rRNA sequences compromises the *in silico* clean-up of remaining rRNA reads from sequencing data, as reads belonging to variable regions would not be removed. To solve this and to enable RNA-seq metagenomics on a broader range of mosquito species, we performed RNA-seq to generate reference rRNA sequences for 33 mosquito species representing 10 genera from Cambodia, the Central African Republic, Madagascar, and French Guiana. Most of these species are associated with vector activity for various pathogens in their respective ecologies (Table 1).

**Figure 1.**
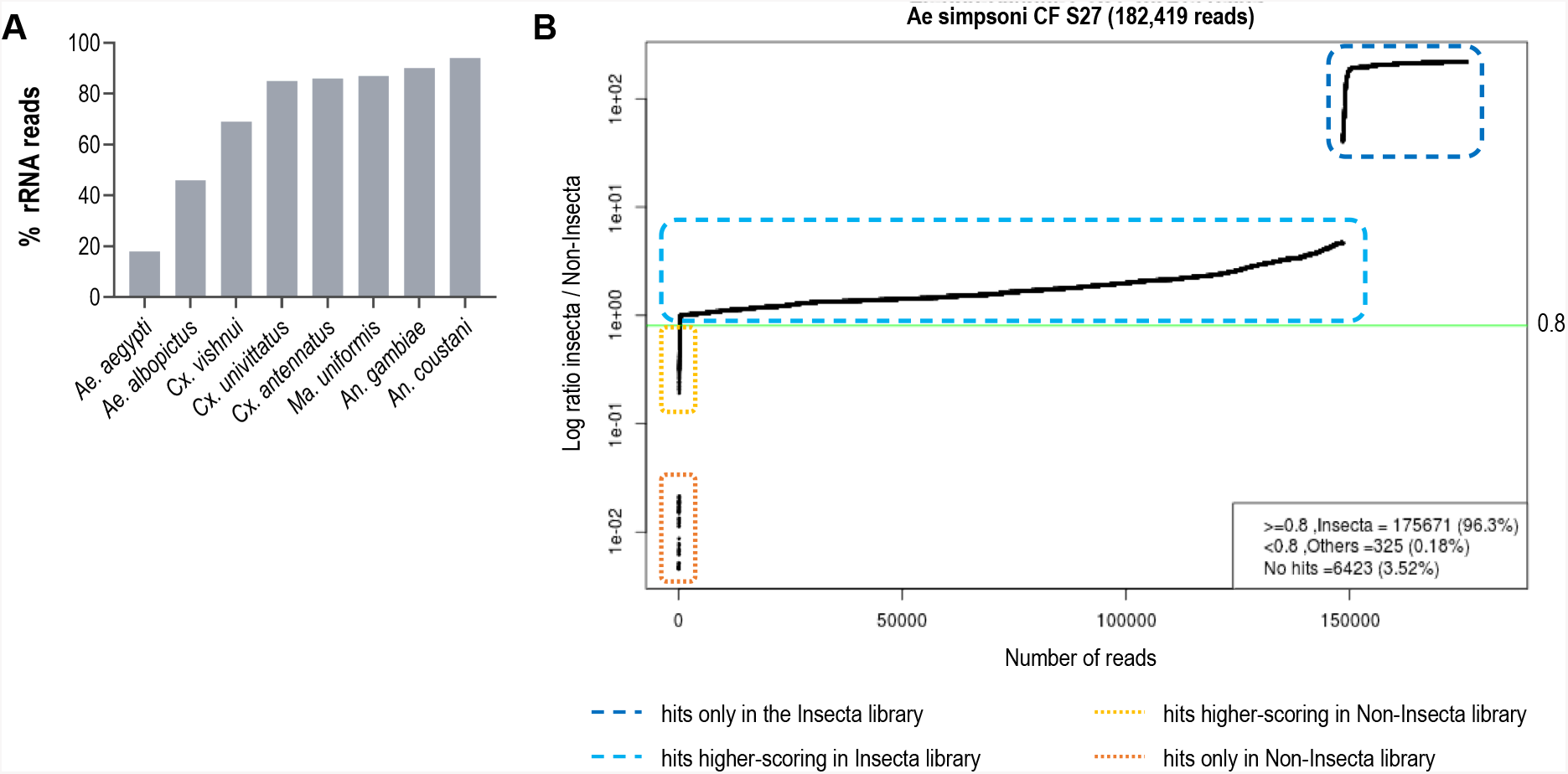
(**A**) Proportion of rRNA reads found in mosquito specimen pools of 5 individuals depleted by probe hybridisation followed by RNase H digestion. Probes were antisense to *Ae. aegypti* rRNA sequences. (**B**) Read vs. score ratio plot of a representative specimen “Ae simpsoni CS S27”. Green line indicates 0.8 cut-off where only reads above this threshold are used in rRNA assembly.

**Table 1.** List of mosquito species represented in this study and their vector status.

| Mosquito taxonomy* | Origin** | Collection site (ecosystem type) | Vector for*** | Reference |
| --- | --- | --- | --- | --- |
| *Aedes (Fredardsius) vittatus* | CF | rural (village) | DENV, ZIKV, CHIKV, YFV | (Diallo et al., 2020) |
| *Aedes (Ochlerotatus) scapularis* | GF | rural (village) | YFV | (Vasconcelos et al., 2001) |
| *Aedes (Ochlerotatus) serratus* | GF | rural (village) | YFV, OROV | (Cardoso et al., 2010; Gaillet et al., 2021) |
| *Aedes (Stegomyia) aegypti* | CF | urban | DENV, ZIKV, CHIKV, YFV | (Kraemer et al., 2019) |
| *Aedes (Stegomyia) albopictus* | CF, KH | rural (village, nature reserve) | DENV, ZIKV, CHIKV, YFV, JEV | (Auerswald et al., 2021; Kraemer et al., 2019) |
| *Aedes (Stegomyia) simpsoni* | CF | rural (village) | YFV | (Mukwaya et al., 2000) |
| *Anopheles (Anopheles) baezai* | KH | rural (nature reserve) | unreported | – |
| *Anopheles (Anopheles) coustani* | MG, CF | rural (village) | RVFV, malaria | (Mwangangi et al., 2013; Nepomichene et al., 2018; Ratovonjato et al., 2011) |
| *Anopheles (Cellia) funestus* | MG, CF | rural (village) | ONNV, malaria | (Lutomiah et al., 2013) |
| *Anopheles (Cellia) gambiae* | MG, CF | rural (village) | ONNV, malaria | (Braut et al., 2004) |
| *Anopheles (Cellia) squamosus* | MG | rural (village) | RVFV, malaria | (Ratovonjato et al., 2011; Stevenson et al., 2016) |
| *Coquillettidia (Rhynchotaenia) venezuelensis* | GF | rural (village) | OROV | (Gaillet et al., 2021) |
| *Culex (Culex) antennatus* | MG | rural (village) | RVFV | (Nepomichene et al., 2018; Ratovonjato et al., 2011) |
| *Culex (Culex) duttoni* | CF | rural (village) | unreported | – |
| *Culex (Culex) neavei* | MG | rural (village) | USUV | (Nikolay et al., 2011) |
| *Culex (Culex) orientalis* | KH | rural (nature reserve) | JEV | (Kim et al., 2015) |
| *Culex (Culex) perexiguus* | MG | rural (village) | WNV | (Vázquez González et al., 2011) |
| *Culex (Culex) pseudovishnui* | KH | rural (nature reserve) | JEV | (Auerswald et al., 2021; Maquart et al., 2021) |
| *Culex (Culex) quinquefasciatus* | MG, CF, KH | rural (village, nature reserve) | ZIKV, JEV, WNV, DENV, SLEV, RVFV, *Wuchereria bancrofti* | (Bhattacharya et al., 2016; Maquart et al., 2021; Ndiaye et al., 2016) |
| *Culex (Culex) tritaeniorhynchus* | MG, KH | rural (village, nature reserve) | JEV, WNV, RVFV | (Auerswald et al., 2021; Maquart et al., 2021) |
| *Culex (Melanoconion) spissipes* | GF | rural (village) | VEEV | (Weaver et al., 2004) |
| *Culex (Melanoconion) portesi* | GF | rural (village) | VEEV, TONV | (Talaga et al., 2021; Weaver et al., 2004) |
| *Culex (Melanoconion) pedroi* | GF | rural (village) | EEEV, VEEV, MADV | (Talaga et al., 2021; M. J. Turell et al., 2008) |
| *Culex (Oculeomyia) bitaeniorhynchus* | MG, KH | rural (village, nature reserve) | JEV | (Auerswald et al., 2021; Maquart et al., 2021) |
| *Culex (Oculeomyia) poicilipes* | MG | rural (village) | RVFV | (Ndiaye et al., 2016) |
| *Eretmapodites intermedius* | CF | rural (village) | unreported | – |
| *Limatus durhamii* | GF | rural (village) | ZIKV | (Barrio-Nuevo et al., 2020) |
| *Mansonia (Mansonia) titillans* | GF | rural (village) | VEEV, SLEV | (Hoyos-López et al., 2015; Michael J. Turell, 1999) |
| *Mansonia (Mansonioides) indiana* | KH | rural (nature reserve) | JEV | (Arunachalam et al., 2004) |
| *Mansonia (Mansonioides) uniformis* | MG, CF, KH | rural (village, nature reserve) | WNV, RVFV, *Wuchereria bancrofti* | (Lutomiah et al., 2013; Maquart et al., 2021; Ughasi et al., 2012) |
| *Mimomyia (Etorleptomyia) mediolineata* | MG | rural (village) | unreported | – |
| *Psorophora (Janthinosoma) ferox* | GF | rural (village) | ROCV | (Mitchell et al., 1986) |
| *Uranotaenia (Uranotaenia) geometrica* | GF | rural (village) | unreported | – |

\* () indicates subgenus
\*\* Origin countries are listed as their ISO alpha-2 codes: Central African Republic, CF; Cambodia, KH;
Madagascar, MG; French Guiana, GF.
\*\* dengue virus, DENV; Zika virus, ZIKV; chikungunya virus, CHIKV; Yellow Fever virus, YFV; Oropouche virus, OROV; Japanese encephalitis virus, JEV; Rift Valley Fever virus, RVFV; O’Nyong Nyong virus, ONNV; Usutu virus, USUV; West Nile virus, WNV; Saint Louis encephalitis virus, SLEV; Venezuelan equine encephalitis virus, VEEV; Tonate virus, TONV; Eastern equine encephalitis virus, EEEV; Madariaga virus, MADV; Rocio virus, ROCV.

### rRNA reads filtering and sequence assembly

Assembling Illumina reads to reconstruct rRNA sequences from total mosquito RNA is not a straightforward task. Apart from host rRNA, total RNA samples also contain rRNA from other organisms associated with the host (microbiota, external parasites, or ingested diet). As rRNA sequences share high homology in conserved regions, Illumina reads (150 bp) from non-host rRNA can interfere with the contig assembly of host 28S and 18S rRNA.

Our score-based filtration strategy, described in detail in Methods, allowed us to bioinformatically remove interfering rRNA reads and achieve successful *de novo* assembly of 28S and 18S rRNA sequences for all our specimens. Briefly, for each Illumina read, we computed a ratio of BLAST scores against an Insecta library over a Non-Insecta library. Reads were segregated into four categories: (i) reads mapping only to the Insecta library, (ii) reads mapping better to the Insecta relative to Non-Insecta library, (iii) reads mapping better to the Non-Insecta relative to the Insecta library, and finally (iv) reads mapping only to the Non-Insecta library (Figure 1B, Supplementary Figure S1). By applying a conservative threshold of 0.8 to account for the non-exhaustiveness of the SILVA database, we removed reads that likely do not originate from mosquito rRNA. Notably, 15 of our specimens were engorged with vertebrate blood, a rich source of non-mosquito rRNA (Supplementary Table S1). The successful assembly of complete 28S and 18S rRNA sequences demonstrates that this strategy performs as expected even with high amounts of non-host rRNA reads. This is particularly important in studies on field-captured mosquitoes as females are often sampled already having imbibed a blood meal or captured using the human landing catch technique.

We encountered challenges for three specimens morphologically identified as *Ma. africana* (Specimen ID 33-35). COI amplification by PCR did not produce any product, hence COI barcoding could not be used to confirm species identity. In addition, SPAdes was only able to assemble partial length contigs, despite the high number of reads with high scores against the Insecta library. Among other *Mansonia* specimens, the partial length contigs shared the highest similarity with contigs obtained from sample “Ma uniformis CF S51”. We then performed a guided assembly using the 28S and 18S sequences of this specimen as references, which successfully produced full-length contigs. In two of these specimens (Specimen ID 34 and 35), our assembly initially produced two sets of 28S and 18S rRNA sequences, one of which was similar to mosquito rRNA with low coverage and another with ten-fold higher coverage and 95% nucleotide sequence similarity to a water mite of genus *Horreolanus* known to parasitize mosquitoes. Our success in obtaining rRNA sequences for mosquito and water mite shows that our strategy can be applied to metabarcoding studies where the input material comprises multiple insect species, provided that appropriate reference sequences of the target species or of a close relative are available.

Altogether, we were able to assemble 122 28S and 114 18S full-length rRNA sequences for 33 mosquito species representing 10 genera sampled from four countries across three continents. This dataset contains, to our knowledge, the first records for 30 mosquito species and for seven genera: *Coquillettidia, Mansonia, Limatus, Mimomyia, Uranotaenia, Psorophora*, and *Eretmapodites*. Individual GenBank accession numbers for these sequences and specimen information are listed in Supplementary Table S1.

### Comparative phylogeny of novel rRNA sequences relative to existing records

To verify the assembly accuracy of our rRNA sequences, we constructed a comprehensive phylogenetic tree from the 28S rRNA sequences generated from our study alongside those publicly available from GenBank (Figure 2). We applied a search criterion for GenBank sequences with at least 95% coverage of our sequence lengths (∼4000 bp), aiming to represent as many species or genera as possible. Although we rarely found records for the same species included in our study, the resulting tree showed that our 28S sequences generally clustered according to their respective species and subgenera, supported by moderate to good bootstrap support at terminal nodes. Species taxa generally formed monophyletic clades, with the exception of *An. gambiae* and *Cx. quinquefasciatus. An. gambiae* 28S rRNA sequences formed a clade with closely related sequences from *An. arabiensis, An. merus*, and *An. coluzzii*, suggesting unusually high interspecies homology for Anophelines or other members of subgenus *Cellia*. Meanwhile, *Cx. quinquefasciatus* 28S rRNA sequences formed a taxon paraphyletic to sister species *Cx. pipiens*.

**Figure 2.**
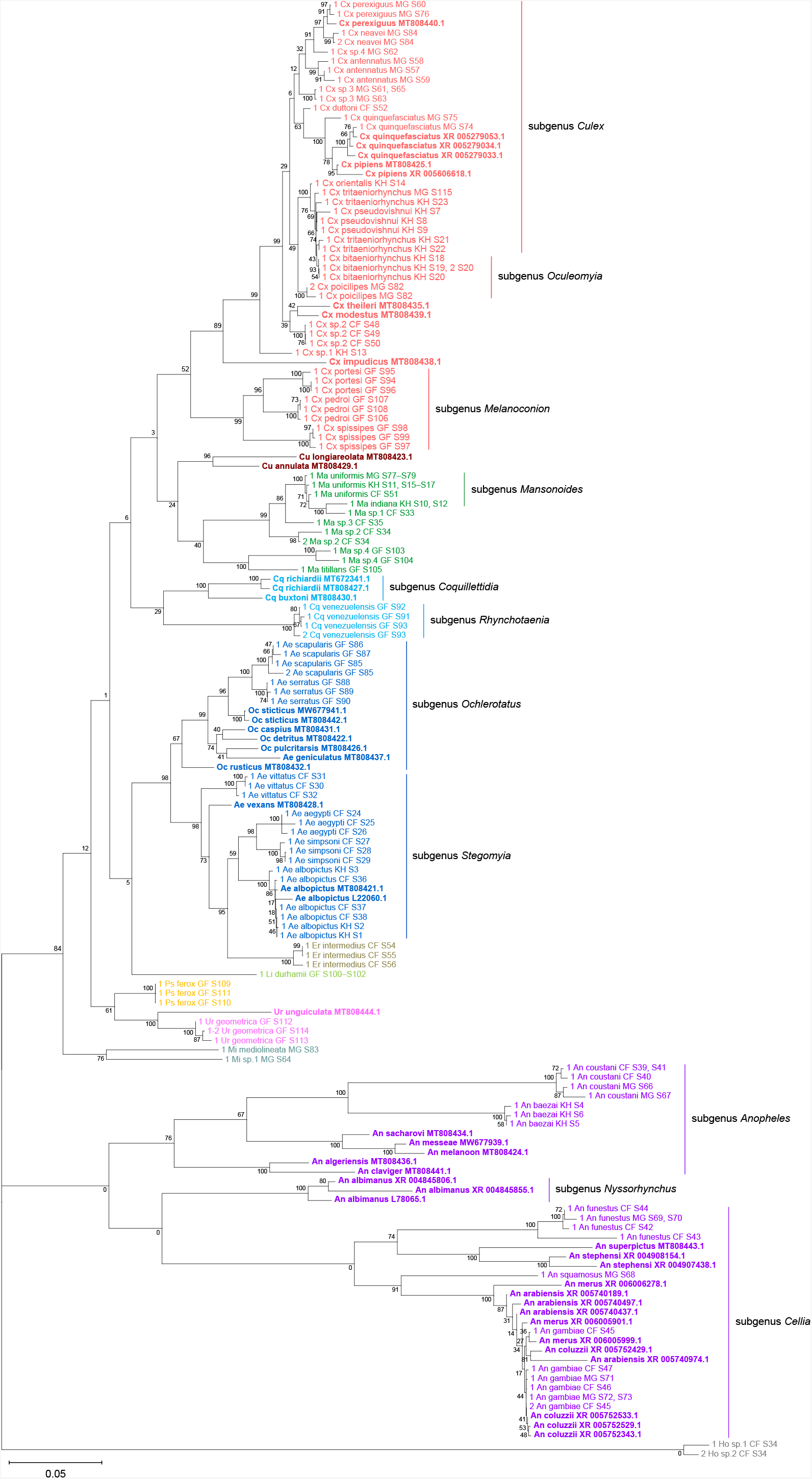
Phylogenetic tree based on 28S sequences generated from this study and from GenBank (3900 bp) as inferred using maximum-likelihood method and constructed to scale in MEGA X (S. Kumar et al., 2018). Values at each node indicate bootstrap support (%) from 500 replications. For sequences from this study, each specimen label contains information on taxonomy, origin (in 2-letter country codes), and specimen ID. Labels in bold indicate GenBank sequences with accession numbers shown. Label colours indicate genera: *Culex* in coral, *Anopheles* in purple, *Aedes* in dark blue, *Mansonia* in dark green, *Culiseta* in maroon, *Limatus* in light green, *Coquillettidia* in light blue, *Psorophora* in yellow, *Mimomyia* in teal, *Uranotaenia* in pink and *Eretmapodites* in brown. Scale bar at 0.05 is shown.

28S sequence-based phylogenetic reconstructions (Figure 2, with GenBank sequences; Supplementary Figure S3, this study only) showed marked incongruence to that of 18S sequences (Figure 3). Although all rRNA trees show the bifurcation of *Culicidae* into subfamilies *Anophelinae* and *Culicinae*, the recovered intergeneric phylogenetic relationships vary between the 28S and 18S trees and are weakly supported. The 18S tree also exhibited several taxonomic anomalies: (i) the lack of definitive clustering by species within the *Culex* subgenus (ii) the lack of distinction between 18S sequences of *Cx. pseudovishnui* and *Cx. tritaeniorhynchus*; (iii) the placement of Ma sp. 3 CF S35 within a *Culex* clade; and (iv) the lack of a monophyletic *Mimomyia* clade. The topology of the 18S tree suggests higher sequence divergence between the two *Cx. quinquefasciatus* taxa and between the two *Mimomyia* taxa than in the 28S trees. However, 28S and 18S rRNA sequences are encoded by linked loci in rDNA clusters and should not be analysed separately.

**Figure 3.**
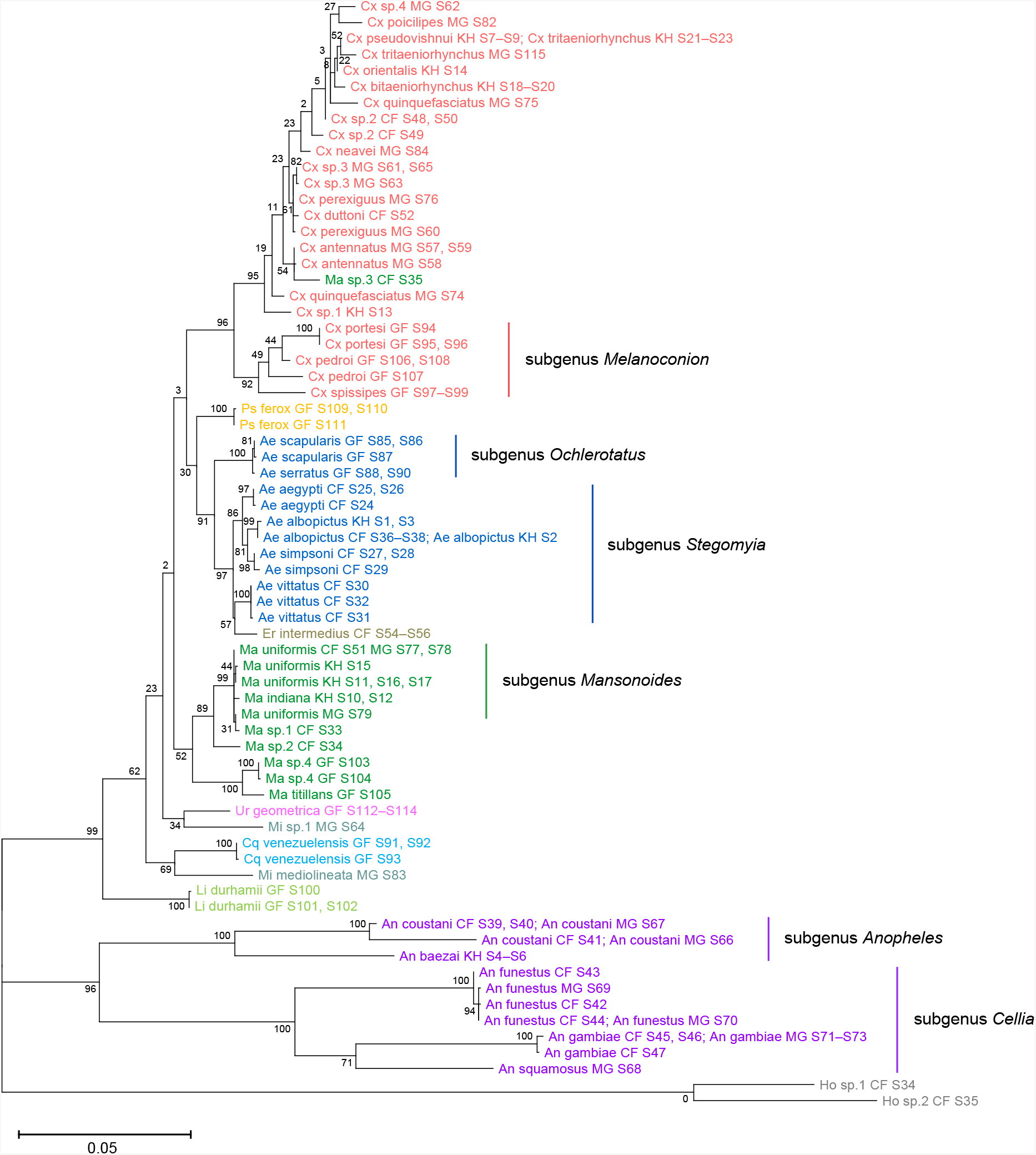
Phylogenetic tree based on 18S sequences (1900 bp) as inferred using maximum-likelihood method and constructed to scale in MEGA X (S. Kumar et al., 2018). Values at each node indicate bootstrap support (%) from 500 replications. Each specimen label contains information on taxonomy, origin (in 2-letter country codes), and specimen ID. Label colours indicate genera: *Culex* in coral, *Anopheles* in purple, *Aedes* in dark blue, *Mansonia* in dark green, *Limatus* in light green, *Coquillettidia* in light blue, *Psorophora* in yellow, *Mimomyia* in teal, *Uranotaenia* in pink and *Eretmapodites* in brown. Scale bar at 0.05 is shown.

Indeed, when concatenated 28S+18S rRNA sequences were generated from the same specimens (Figure 4), the phylogenetic tree resulting from these sequences more closely resembles the 28S tree (Figure 2) with regard to the basal position of the *Mimomyia* clade within the *Culicinae* subfamily with good bootstrap support in either tree (84% in 28S tree, 100% in concatenated 28S+18S tree). For internal nodes, bootstrap support values were higher in the concatenated tree compared to the 28S tree. Interestingly, the 28S+18S tree formed an *Aedini* tribe-clade encompassing taxa from genera *Psorophora, Aedes*, and *Eretmapodites*, possibly driven by the inclusion of 18S sequences. Concatenation also resolved the anomalies found in the 18S tree and added clarity to the close relationship between *Culex* and *Mansonia* taxa. Of note, relative to the 28S tree (Figure 2) the *Culex* and *Mansonia* genera are no longer monophyletic in the concatenated 28S+18S tree (Figure 4). Genus *Culex* is paraphyletic with respect to subgenus *Mansonoides* of genus *Mansonia* (Figure 2). *Ma. titillans* and Ma sp. 4, which we suspect to be *Ma. pseudotitillans*, always formed a distinct branch in 28S or 18S phylogenies, thus possibly representing a clade of subgenus *Mansonia*.

**Figure 4.**
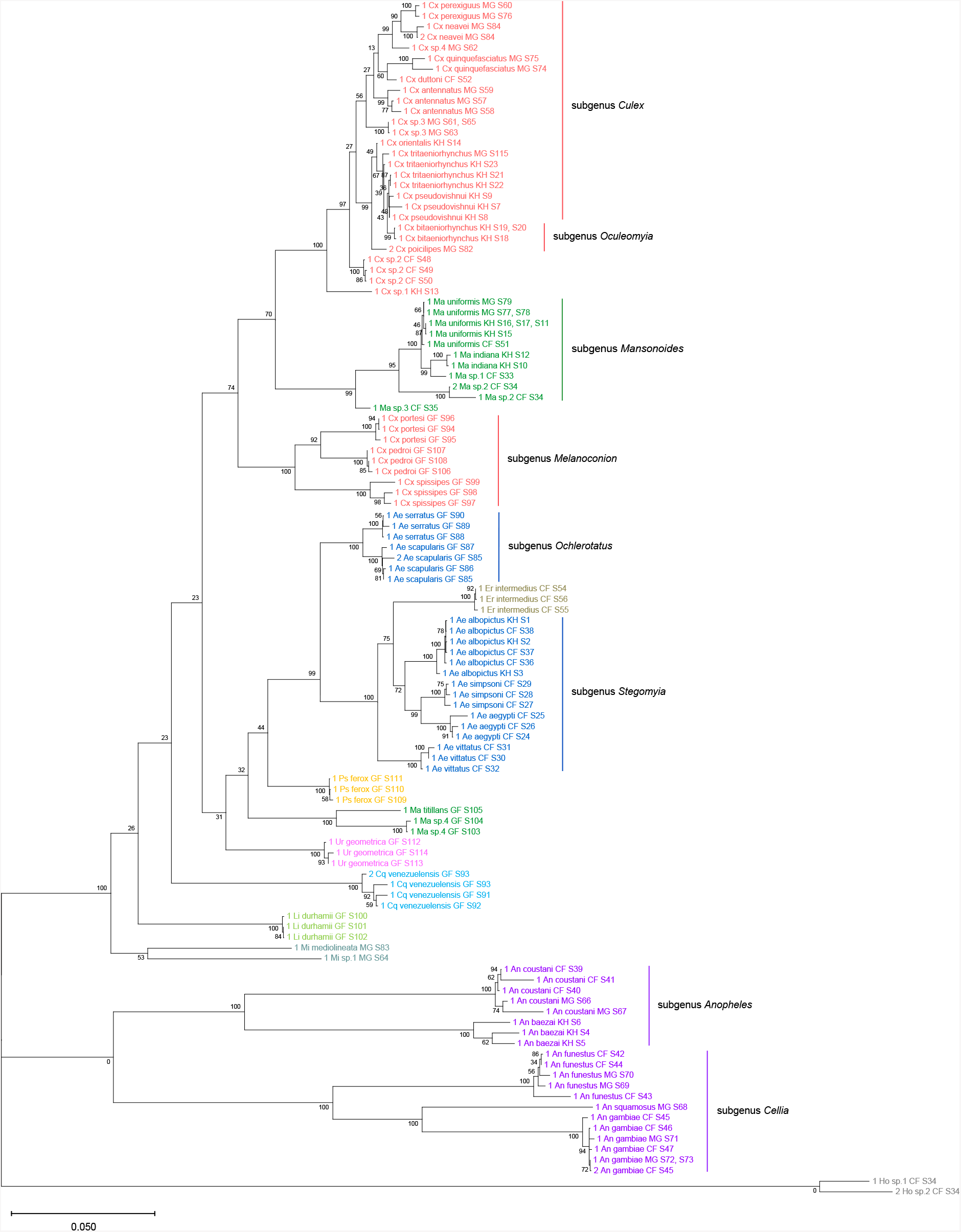
Phylogenetic tree based on concatenated 28S and 18S sequences generated from this study (3900+1900 bp) as inferred using maximum-likelihood method and constructed to scale in MEGA X (S. Kumar et al., 2018). Values at each node indicate bootstrap support (%) from 500 replications. For sequences from this study, each specimen label contains information on taxonomy, origin (as indicated in 2-letter country codes), and specimen ID. Label colours indicate genera: *Culex* in coral, *Anopheles* in purple, *Aedes* in dark blue, *Mansonia* in dark green, *Culiseta* in maroon, *Limatus* in light green, *Coquillettidia* in light blue, *Psorophora* in yellow, *Mimomyia* in teal, *Uranotaenia* in pink and *Eretmapodites* in brown. Scale bar at 0.05 is shown.

The concatenated 28S+18S tree recapitulates what is classically known about the systematics of our specimens, namely (i) the early divergence of subfamily *Anophelinae* from subfamily *Culicinae*, (ii) the division of genus *Anopheles* into two subgenera, *Anopheles* and *Cellia*, (iii) the division of genus *Aedes* into subgenera *Stegomyia* and *Ochlerotatus*, (iv) the divergence of monophyletic subgenus *Melanoconion* within the *Culex* genus (Harbach, 2007; Harbach & Kitching, 2016).

### rRNA as a molecular marker for taxonomy and phylogeny

We sequenced a 621 bp region of the COI gene to confirm morphological species identification of our specimens and to compare the functionality of rRNA and COI sequences as molecular markers for taxonomic and phylogenetic investigations. COI sequences were able to unequivocally determine the species identity in most specimens except for the following cases. *An. coustani* COI sequences from our study regardless of specimen origin shared remarkably high nucleotide similarity (>98%) with several other *Anopheles* species such as *An. rhodesiensis, An. rufipes, An. ziemanni, An. tenebrosus*, although *An. coustani* remained the most frequent and closest match. In the case of *Ae. simpsoni*, three specimens had been morphologically identified as *Ae. opok* although their COI sequences showed 97–100% similarity to that of *Ae. simpsoni*. As GenBank held no records of *Ae. opok* COI at the time of this study, we instead aligned the putative *Ae. simpsoni* COI sequences against two sister species of *Ae. opok*: *Ae. luteocephalus* and *Ae. africanus*. We found they shared only 90% and 89% similarity, respectively. Given this significant divergence, we concluded these specimens to be *Ae. simpsoni*. Ambiguous results were especially frequent among *Culex* specimens belonging to the *Cx. pipiens* or *Cx. vishnui* subgroups, where the query sequence differed with either of the top two hits by a single nucleotide. For example, between *Cx. quinquefasciatus* and *Cx. pipiens* of the *Cx. pipiens* subgroup, and between *Cx. vishnui* and *Cx. tritaeniorhynchus* of the *Cx. vishnui* subgroup.

Among our three specimens of *Ma. titillans*, two appeared to belong to a single species that is different but closely related to *Ma. titillans*. We surmised that these specimens could instead be *Ma. pseudotitillans* based on morphological similarity but were not able to verify this by molecular means as no COI reference sequence is available for this species. These specimens are hence putatively labelled as “Ma sp.4”.

Phylogenetic reconstruction based on the COI sequences showed clustering of all species taxa into distinct clades, underlining the utility of the COI gene in molecular taxonomy (Figure 5)(Hebert et al., 2003; Ratnasingham & Hebert, 2007). However, species delineation among members of *Culex* subgroups were not as clear cut, although sister species were correctly placed as sister taxa. This is comparable to the 28S+18S tree (Figure 4) and is indicative of lower intraspecies distances relative to interspecies distances.

**Figure 5.**
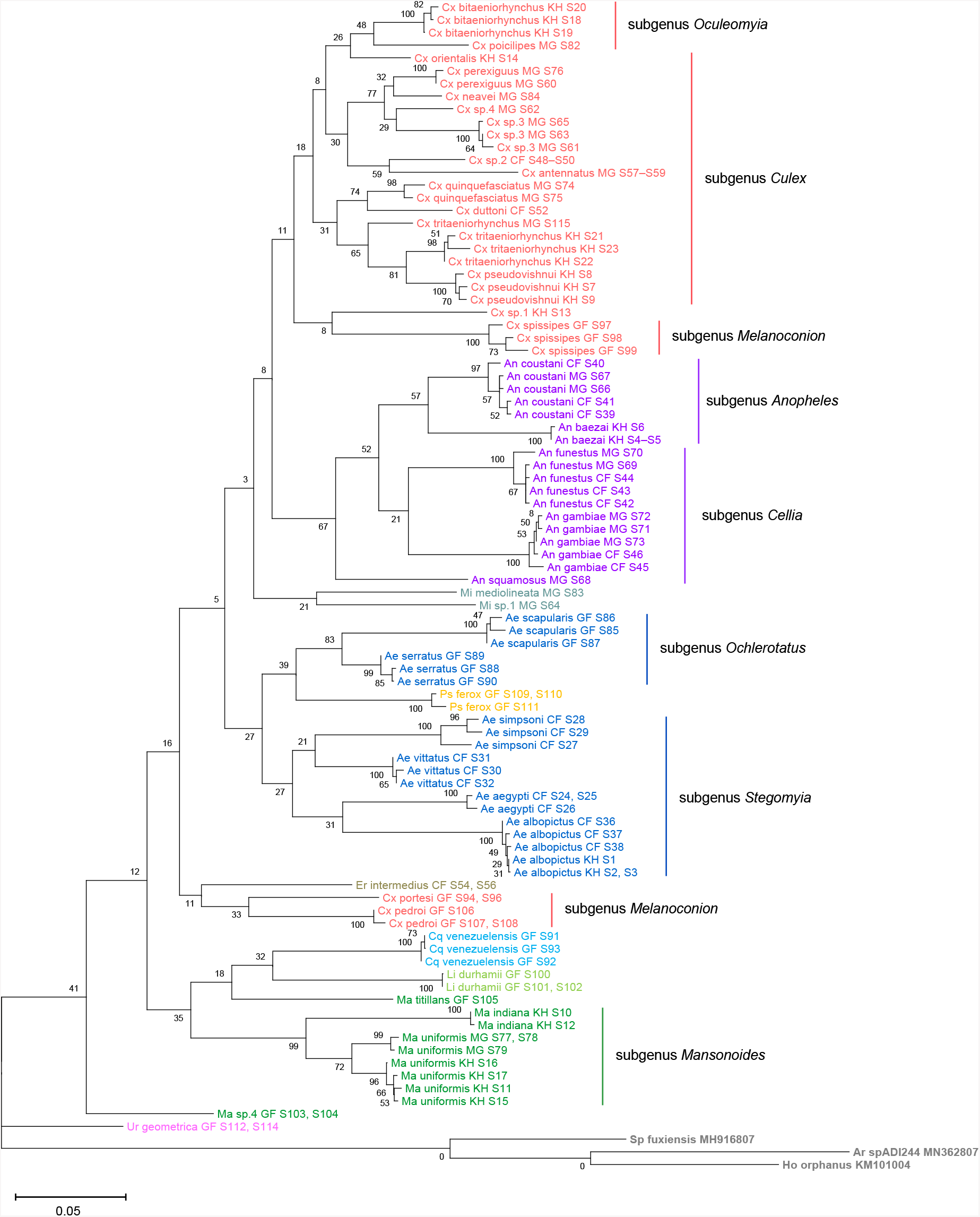
Phylogenetic tree based on COI sequences (621–699 bp) as inferred using maximum-likelihood method and constructed to scale in MEGA X (S. Kumar et al., 2018). Values at each node indicate bootstrap support (%) from 500 replications. Each specimen label contains information on taxonomy, origin (as indicated in 2-letter country codes), and specimen ID. Label colours indicate genera: *Culex* in coral, *Anopheles* in purple, *Aedes* in dark blue, *Mansonia* in dark green, *Limatus* in light green, *Coquillettidia* in light blue, *Psorophora* in yellow, *Mimomyia* in teal, *Uranotaenia* in pink and *Eretmapodites* in brown. Scale bar at 0.05 is shown.

To evaluate the utility of 28S and 18S rRNA sequences for molecular taxonomy, we used the 28S+18S rRNA tree to discern the identity of six specimens for which COI barcoding could not be performed. These specimens include three unknown *Mansonia* species (Specimen ID 33–35), a *Ma. uniformis* (Specimen ID 51), an *An. gambiae* (Specimen ID 47), and a *Ur. geometrica* (Specimen ID 113). Their positions in the 28S+18S rRNA tree relative to adjacent taxa confirms the morphological identification of all six specimens to the genus level and, for three of them, to the species level (Figure 4).

The phylogenetic relationships indicated by the COI tree compared to the 28S+18S tree present only a few points of congruence. COI-based phylogenetic inference indeed showed clustering of generic taxa into monophyletic clades albeit with very weak bootstrap support, except for genera *Culex* and *Mansonia* (Figure 5). Contrary to the 28S+18S tree (Figure 4), *Culex* subgenus *Melanoconion* was depicted as a polyphyletic taxon with *Cx. spissipes* being a part of the greater *Culicini* clade with members from subgenera *Oculeomyia* and *Culex* while *Cx. pedroi* and *Cx. portesi* formed a distantly related clade. Among the *Mansonia* specimens, the two unknown Ma sp.4 specimens were not positioned as the nearest neighbours of *Ma. titillans* and instead appeared to have diverged earlier from most of the other taxa from the *Culicidae* family. Notably, the COI sequences of genus *Anopheles* is not basal to the other members of *Culicidae* and is instead shown to be sister to Culex COI sequences (8% bootstrap support). This is a direct contrast to what is suggested by the rRNA phylogenies (Figures 2–4), which suggests *Culex* rRNA sequences to be among the most recently diverged. Bootstrap support for the more internal nodes of the COI trees were remarkably low compared to those of rRNA-based trees.

In all rRNA trees, it is clear that the interspecific and intersubgeneric evolutionary distances within the genus *Anopheles* are high relative to any other genera, indicating a greater degree of divergence. This is evidenced by the longer branch lengths connecting Anopheline species-clades to the node of the most recent common ancestor for subgenera *Anopheles* and *Cellia* (Figures 2-4, Supplementary Figure S3). This feature is not evident in the COI tree, where the Anopheline interspecies distances are comparable to those within the *Culex, Aedes*, and *Mansonia* taxa.

### On *Culex* subgroups

*Culex* (subgenus *Culex*) specimens of this study comprise several closely related sister species belonging to the *Cx. vishnui* and *Cx. univittatus* subgroups, which are notoriously difficult to differentiate based on morphology. Accordingly, in the 28S+18S rRNA (Figure 4) and COI (Figure 5) trees these species and their known sister species were clustered together within the *Culex* (subgenus *Culex*) clade: *Cx. tritaeniorhynchus* with *Cx. pseudovishnui* (*Cx. vishnui* subgroup); *Cx. perexiguus* with *Cx. neavei* (*Cx. univittatus* subgroup).

The use of COI barcoding to distinguish between members of the *Culex* subgroups was limited. For example, for the two *Cx. quinquefasciatus* samples in our taxonomic assemblage (Specimen ID 74 and 75), BLAST analyses of their COI sequences revealed they are a single nucleotide away from *Cx. pipiens* or *Cx. quinquefasciatus* COI sequences (Supplementary Table S2). In the 28S rRNA tree with GenBank sequences (Figure 2), two GenBank sequences of *Cx. pipiens* sequences formed a clade sister to another containing three *Cx. quinquefasciatus* GenBank sequences and the “Cx quinquefasciatus MG S74” sequence with 78% bootstrap support. This is in accordance with other studies examining mitochondrial sequences (Sun et al., 2019) and morphological attributes (Harbach et al., 2017). This shows that the 28S rRNA sequence can distinguish the two species and confirms that “Cx quinquefasciatus MG S74” is indeed a *Cx. quinquefasciatus* specimen. However, “Cx quinquefasciatus MG S75” is shown to be basal from other sequences within this *Cx. pipiens* subgroup-clade with 100% bootstrap support. Given that *Cx. quinquefasciatus* and *Cx. pipiens* are known to interbreed, it is plausible that this individual is a hybrid of the two species (Farajollahi et al., 2011).

## DISCUSSION

RNA-seq metagenomics on field-captured non-urban mosquitoes is a valuable tool for pre-empting mosquito-borne virus emergences through surveillance and virus discovery. However, the lack of reference rRNA sequences hinders good oligo-based depletion and efficient clean-up of RNA-seq data. Additionally, *de novo* assembly of rRNA sequences is complicated due to regions that are highly conserved across all distantly related organisms that could be present in a single specimen, i.e., microbiota, parasites, or vertebrate blood meal. Hence, we sought to establish a method to bioinformatically filter out non-host rRNA reads for the accurate assembly of novel 28S and 18S rRNA reference sequences.

We found that phylogenetic reconstructions based on 28S sequences or concatenated 28S+18S rRNA sequences were able to correctly cluster mosquito taxa according to species and corroborate current mosquito classification. This demonstrates that our bioinformatics methodology reliably generates bona fide 28S and 18S rRNA sequences, even in specimens parasitized by water mites or engorged with vertebrate blood. Further, we were able to use 28S+18S rRNA taxonomy for molecular species identification when COI sequences were unavailable or ambiguous, thus supporting the use of rRNA sequences as a barcode. They have the advantage of circumventing the need to additionally isolate and sequence DNA from specimens, as RNA-seq reads can be directly mapped against reference sequences. Post-depletion, in our hands there are sufficient numbers of remaining reads (5–10% of reads per sample) for assembly of complete rRNA contigs (unpublished data).

Phylogenetic inferences based on 28S or 18S rRNA sequences alone do not recover the same interspecific relationships. Relative to 28S sequences, we observed more instances where multiple specimens have near-identical 18S rRNA sequences. This can occur for specimens belonging to the same species, but also for conspecifics sampled from different geographic locations, such as *An. coustani, An. gambiae*, or *Ae. albopictus*. More rarely, specimens from the same species subgroup, such as *Cx. pseudovishnui* and *Cx. tritaeniorhynchus*, also shared 18S rRNA sequences. This was surprising given that the 18S rRNA sequences in our dataset is 1900 bp long. Concatenation of 28S and 18S rRNA sequences resolved this issue, enabling species delineation even among sister species of *Culex* subgroups, with which morphological identification meets its limits.

In Cambodia and other parts of Asia, the *Cx. vishnui* subgroup includes *Cx. tritaeniorhynchus, Cx. vishnui*, and *Cx. pseudovishnui*, which are important vectors of JEV (Maquart & Boyer, 2022). The former two were morphologically identified in our study but later revealed by COI barcoding to be a sister species. Discerning sister species of the *Cx. pipiens* subgroup is further complicated by interspecific breeding, with some populations showing genetic introgression to varying extents (Cornel et al., 2003). The seven sister species of this subgroup are practically indistinguishable based on morphology and require molecular methods to discern (Farajollahi et al., 2011; Zittra et al., 2016). Indeed, the 621 bp COI sequence amplified in our study did not contain enough nucleotide divergence to allow clear identification, given that the COI sequence of *Cx. quinquefasciatus* specimens differed from that of *Cx. pipiens* by a single nucleotide. Batovska et al. (2017) found that even the Internal Transcribed Spacer 2 (ITS2) rDNA region, another common molecular marker, could not differentiate the two species. Other DNA molecular markers such as nuclear Ace-2 or CQ11 genes (Aspen & Savage, 2003; Zittra et al., 2016) or *Wolbachia pipientis* infection status (Cornel et al., 2003) are typically employed in tandem. In our study, 28S rRNA sequence-based phylogeny (Figure 2) validated the identity of specimen “Cx quinquefasciatus MG S74” and suggested that specimen “Cx quinquefasciatus MG S75” might have been a *pipiens*-*quinquefasciatus* hybrid. These examples demonstrate how 28S rRNA sequences, concatenated with 18S or alone, contain enough resolution to differentiate between *Cx. pipiens* and *Cx. quinquefasciatus*. rRNA-based phylogeny thus allows for more accurate species identification and ecological observations in the context of disease transmission. Additionally, tracing the genetic flow across hybrid populations within the *Cx. pipiens* subgroup can inform estimates of vectorial capacity for each species. As only one or two members from the *Cx. pipiens* and *Cx. vishnui* subgroups were represented in our taxonomic assemblage, an explicit investigation including all member species of these subgroups in greater sample numbers is warranted to further test the degree of accuracy with which 28S and 18S rRNA sequences can delineate sister species.

Our study included French Guianese *Culex* species *Cx. spissipes* (group Spissipes*), Cx. pedroi* (group Pedroi), and *Cx. portesi* (group Vomerifer). These species belong to the New World subgenus *Melanoconion*, section Spissipes, with well-documented distribution in North and South Americas (Sirivanakarn, 1982) and are vectors of encephalitic alphaviruses EEEV and VEEV among others (Talaga et al., 2021; M. J. Turell et al., 2008; Weaver et al., 2004). Indeed, our rooted rRNA and COI trees showed the divergence of the three *Melanoconion* species from the major *Culex* clade comprising species broadly found across Africa and Asia (Auerswald et al., 2021; Farajollahi et al., 2011; Nchoutpouen et al., 2019; Takhampunya et al., 2011). The topology of the concatenated 28S+18S tree places the *Cx. portesi* and *Cx. pedroi* species-clades as sister groups (92% bootstrap support), with *Cx. spissipes* as a basal group within the *Melanoconion* clade (100% bootstrap support). This corroborates the systematics elucidated by Navarro and Weaver (2004) using the ITS2 marker, and those by Sirivanakarn (1982) and Sallum and Forattini (1996) based on morphology. Curiously, in the COI tree, *Cx. spissipes* sequences were clustered with unknown species Cx. sp1, forming a clade sister to another containing other *Culex* (*Culex*) and *Culex* (*Oculeomyia*) species, albeit with very low bootstrap support. Previous phylogenetic studies based on the COI gene have consistently placed *Cx. spissipes* or the Spissipes group basal to other groups within the *Melanoconion* subgenus (Torres-Gutierrez et al., 2016, 2018). However, these studies contain only *Culex* (*Melanoconion*) species in their assemblage, apart from *Cx. quinquefasciatus* to act as an outgroup. This clustering of *Cx. spissipes* with non-*Melanoconion* species in our COI phylogeny could be an artefact of a much more diversified assemblage rather than a true phylogenetic link.

Taking advantage of our multi-country sampling, we examined whether rRNA or COI phylogeny can be used to discriminate conspecifics originating from different geographies. Our assemblage contains five of such species: *An. coustani, An. funestus, An. gambiae, Ae. albopictus*, and *Ma. uniformis*. Among the rRNA trees, the concatenated 28S+18S tree and 28S tree were able to discriminate between *Ma. uniformis* specimens from Madagascar, Cambodia, and the Central African Republic, and between *An. coustani* specimens from Madagascar and the Central African Republic (100% bootstrap support). In the COI tree, only *Ma. uniformis* was resolved into geographical clades comprising specimens from Madagascar and specimens from Cambodia (72% bootstrap support). No COI sequence was obtained from one *Ma. uniformis* from the Central African Republic. The 28S+18S rRNA sequences ostensibly provided more population-level genetic information than COI sequences alone with better support. The use of rRNA sequences in investigating the biodiversity of mosquitoes should therefore be explored with a more comprehensive taxonomic assemblage.

The phylogenetic reconstructions based on rRNA or COI sequences in our study are hardly congruent, but two principal differences stand out. First, the COI phylogeny does not recapitulate the early divergence of *Anophelinae* from *Culicinae* (Figure 5). This is at odds with other studies estimating mosquito divergence times based on mitochondrial genes (Logue et al., 2013; Lorenz et al., 2021) or nuclear genes (Reidenbach et al., 2009). The second notable feature in the rRNA trees is the remarkably large interspecies and intersubgeneric evolutionary distances within genus *Anopheles* relative to *Culicinae* genera (Figures 2–4, Supplementary Figure S4) but this is not apparent in the COI tree. The hyperdiversity among *Anopheles* taxa may be attributed to the earlier diversification of the *Anophelinae* subfamily in the early Cretaceous period compared to that of the *Culicinae* subfamily, a difference of at least 40 million years (Lorenz et al., 2021). The differences in rRNA and COI tree topologies indicate a limitation in using COI alone to determine evolutionary relationships. Importantly, drawing phylogenetic conclusions from short DNA barcodes such as COI has been cautioned against due to its weak phylogenetic signal (Hajibabaei et al., 2006). The relatively short length of our COI sequences (621–699 bp) combined with the 100-fold higher nuclear substitution rate of mitochondrial genomes relative to nuclear genomes (Arctander, 1995) could result in homoplasy (Danforth et al., 2005), making it difficult to clearly discern ancestral sequences and correctly assign branches into lineages, as evidenced by the poor nodal bootstrap support at genus-level branches. Indeed, in the study by Lorenz et al. (2021), a phylogenetic tree constructed using a concatenation of all 13 protein-coding genes of the mitochondrial genome was able to resolve ancient divergence events. This affirms that while COI sequences can be used to reveal recent speciation events, longer or multi-gene molecular markers are necessary for studies into deeper evolutionary relationships (Danforth et al., 2005).

In contrast to Anophelines where 28S rRNA phylogenies illustrated higher interspecies divergence compared to COI phylogeny, two specimens of an unknown *Mansonia* species, “Ma sp.4 GF S103” and “Ma sp.4 GF S104”, provided an example where interspecies relatedness based on their COI sequences is greater than that based on their rRNA sequences in relation to “Ma titillans GF S105”. While all rRNA trees (Figure 2–4) placed “Ma titillans GF S105” as a sister taxon with 100% bootstrap support, the COI tree placed Ma sp.4 basal to all other species except *Ur. geometrica*. This may hint at a historical selective sweep in the mitochondrial genome, whether arising from geographical separation, mutations, or linkage disequilibrium with inherited symbionts (Hurst & Jiggins, 2005), resulting in the disparate mitochondrial haplogroups found in French Guyanese Ma sp.4 and *Ma. titillans*. In addition, both haplogroups are distant from those associated with members of subgenus *Mansonoides*. To note, the COI sequences of “Ma sp.4 GF S103” and “Ma sp.4 GF S104” share 87.12 and 87.39% nucleotide similarity, respectively, to that of “Ma titillans GF S105”. Interestingly, the endosymbiont *Wolbachia pipientis* has been detected in *Ma. titillans* sampled from Brazil (De Oliveira et al., 2015), which may contribute to the divergence of “Ma titillans GF S105” COI sequence away from those of Ma sp.4. This highlights other caveats of using a mitochondrial DNA marker in determining evolutionary relationships (Hurst & Jiggins, 2005), which nuclear markers such as 28S and 18S rRNA sequences may be immune to.

## Conclusions

RNA-seq metagenomics is a valuable tool for surveillance and virus discovery in non-urban mosquitoes but it is impeded by the lack of full-length rRNA reference sequences. Here we presented a rRNA sequence assembly strategy and 234 newly generated 28S and 18S mosquito rRNA sequences. Our work has expanded the current rRNA reference library by providing, to our knowledge, the first full-length rRNA records for 30 species in public databases and paves the way for the assembly of many more. These novel rRNA sequences can improve mosquito RNA-seq metagenomics by expanding reference sequence data for the optimization of species-specific oligo-based rRNA depletion protocols, for streamlined species identification by rRNA barcoding and for improved RNA-seq data clean-up. In addition, RNA barcodes could serve as an additional tool for mosquito taxonomy and phylogeny although further studies are necessary to reveal how they compare against other nuclear or mitochondrial DNA marker systems (Batovska et al., 2017; Beebe, 2018; Behura, 2006; Ratnasingham & Hebert, 2007; Reidenbach et al., 2009; Vezenegho et al., 2022).

We showed that phylogenetic inferences from a tree based on 28S rRNA sequences alone or concatenated 28S +18S rRNA sequences largely agree with contemporary mosquito classification and can be used for species diagnostics given a reference sequence. In analysing the same set of specimens by COI or rRNA sequences, we found deep discrepancies in phylogenetic inferences. We conclude that while COI-based phylogeny can reveal recent speciation events, rRNA sequences may be better suited for investigations of deeper evolutionary relationships as they are less prone to selective sweeps and homoplasy.

## MATERIALS AND METHODS

### Sample collection

Mosquito specimens were sampled from 2019 to 2020 by medical entomology teams from the Institut Pasteur de Bangui (Central African Republic, Africa; CF), Institut Pasteur de Madagascar (Madagascar, Africa; MG), Institut Pasteur du Cambodge (Cambodia, Asia; KH), and Institut Pasteur de la Guyane (French Guiana, South America; GF). Adult mosquitoes were sampled using several techniques including CDC light traps, BG sentinels, and human-landing catches. Sampling sites are non-urban locations including rural settlements in the Central African Republic, Madagascar, and French Guiana and national parks in Cambodia. Mosquitoes were identified using morphological identification keys on cold tables before preservation by flash freezing in liquid nitrogen and transportation in dry ice to Institut Pasteur Paris for analysis. A list of the 112 mosquito specimens included in our taxonomic assemblage and their related information are provided in Supplementary Table S1.

### RNA and DNA isolation

Nucleic acids were isolated from mosquito specimens using TRIzol reagent according to manufacturer’s protocol (Invitrogen, Thermo Fisher Scientific, Waltham, Massachusetts, USA). Single mosquitoes were homogenised into 200 µL of TRIzol reagent and other of the reagents within the protocol were volume-adjusted accordingly. Following phase separation, RNA were isolated from the aqueous phase while DNA were isolated from the remaining interphase and phenol-chloroform phase. From here, RNA is used to prepare cDNA libraries for next generation sequencing while DNA is used in PCR amplification and Sanger sequencing of the mitochondrial *cytochrome* c *oxidase subunit I* (COI) gene as further described below.

### Probe depletion of rRNA

We tested a selective rRNA depletion protocol by Morlan *et al*. (2012) on several mosquito species from the *Aedes, Culex*, and *Anopheles* genera. We designed 77 tiled 80 bp DNA probes antisense to the *Ae. aegypti* 28S, 18S, and 5.8S rRNA sequences. A pool of probes at a concentration of 0.04 µM were prepared. To bind probes to rRNA, 1 µL of probes and 2 µL of Hybridisation Buffer (100 mM Tris-HCl and 200 mM NaCl) was added to rRNA samples to a final volume of 20 µL and subjected to a slow-cool incubation starting at 95 °C for 2 minutes, then cooling to 22 °C at a rate of 0.1 °C per second, ending with an additional 5 minutes at 22 °C. The resulting RNA:DNA hybrids were treated with 2.5 µL Hybridase™ Thermostable RNase H (Epicentre, Illumina, Madison, Wisconsin, USA) and incubated at 37 °C for 30 minutes. To remove DNA probes, the mix was treated with 1 µL DNase I (Invitrogen) and purified with Agencourt RNAClean XP Beads (Beckman Coulter, Brea, California, USA). The resulting RNA is used for total RNA sequencing to check depletion efficiency.

### Total RNA sequencing

To obtain rRNA sequences, RNA samples were quantified on a Qubit Fluorometer (Invitrogen) using the Qubit RNA BR Assay kit (Invitrogen) for concentration adjustment. Non-depleted total RNA was used for library preparation for next generation sequencing using the NEBNext Ultra II RNA Library Preparation Kit for Illumina (New England Biolabs, Ipswich, Massachusetts, USA) and the NEBNext Multiplex Oligos for Illumina (Dual Index Primers Set 1) (New England Biolabs). Sequencing was performed on a NextSeq500 sequencing system (Illumina, San Diego, California, USA). Quality control of fastq data and trimming of adapters were performed with FastQC and cutadapt, respectively.

### 28S and 18S rRNA assembly

To obtain 28S and 18S rRNA contigs, we had to first clean our fastq library by separating the reads representing mosquito rRNA from all other reads. To achieve this, we used the SILVA RNA sequence database to create 2 libraries: one containing all rRNA sequences recorded under the “Insecta” node of the taxonomic tree, the other containing the rRNA sequences of many others nodes distributed throughout the taxonomic tree, hence named “Non-Insecta” (Quast et al., 2013). Each read was aligned using the nucleotide Basic Local Alignment Search Tool (BLASTn, **https://blast.ncbi.nlm.nih.gov/**) of the National Center for Biotechnology Information (NCBI) against each of the two libraries and the scores of the best high-scoring segment pairs from the two BLASTns are subsequently used to calculate a ratio of Insecta over Non-Insecta scores (Altschul et al., 1990). Only reads with a ratio greater than 0.8 were used in the assembly. The two libraries being non-exhaustive, we chose this threshold of 0.8 to eliminate only reads that were clearly of a non-insect origin. Selected reads were assembled with the SPAdes assembler using the “-rna” option, allowing more heterogeneous coverage of contigs and kmer lengths of 31, 51 and 71 bases (Bankevich et al., 2012). This method successfully assembled rRNA sequences for all specimens, including a parasitic *Horreolanus* water mite (122 sequences for 28S and 114 sequences for 18S).

Initially, our filtration technique had two weaknesses. First, there is a relatively small number of complete rRNA sequences in the Insecta library from SILVA. To compensate for this, we carried out several filtration cycles, each time adding in the complete sequences produced in previous cycles to the Insecta library. Second, when our mosquito specimens were parasitized by other insects, it was not possible to bioinformatically filter out rRNA reads belonging to the parasite. For these rare cases, we used the “--trusted-contigs” option of SPAdes, giving it access to the 28S and 18S sequences of the mosquito closest in terms of taxonomic distance. By doing this, the assembler was able to reconstruct the rRNA of the mosquito as well as the rRNA of the parasitizing insect. All assembled rRNA sequences from this study have been deposited in GenBank with accession numbers OM350214–OM350327 for 18S rRNA sequences and OM542339–OM542460 for 28S rRNA sequences.

### COI amplicon sequencing

The mitochondrial COI gene was amplified from DNA samples using the universal “Folmer” primer set LCO1490 (5’-GGTCAACAAATCATAAAGATATTGG -3’) and HCO2198 (5’-TAAACTTCAGGGTGACCAAAAAATCA-3’), as per standard COI barcoding practices, producing a 658 bp product (Folmer et al., 1994). PCRs were performed using Phusion High-Fidelity DNA Polymerase (Thermo Fisher Scientific). Every 50 µL reaction contained 10 µL of 5X High Fidelity buffer, 1 µL of 10 mM dNTPs, 2.5 µL each of 10 mM forward (LCO1490) and reverse (HCO2198) primer, 28.5 µL of water, 5 µL of DNA sample, and 0.5 µL of 2 U/µL Phusion DNA polymerase. A 3-step cycling incubation protocol was used: 98 °C for 30 seconds; 35 cycles of 98 °C for 10 seconds, 60 °C for 30 seconds, and 72 °C for 15 seconds; 72 °C for 5 minutes ending with a 4 °C hold. PCR products were size-verified using gel electrophoresis and then gel-purified using the QIAquick Gel Extraction Kit (Qiagen, Hilden, Germany). Sanger sequencing of the COI amplicons were performed by Eurofins Genomics, Ebersberg, Germany.

### COI sequence analysis

Forward and reverse COI DNA sequences were end-trimmed to remove bases of poor quality (Q score < 30). At the 5’ ends, sequences were trimmed at the same positions such that all forward sequences start with 5’-TTTTGG and all reverse sequences start with 5’-GGNTCT. Forward and reverse sequences were aligned using BLAST to produce a 621 bp consensus sequence. In cases where good quality sequences extends beyond 621 bp, forward and reverse sequences were assembled using Pearl (**https://www.gear-genomics.com/pearl/**) and manually checked for errors against trace files (Rausch et al., 2019, 2020). We successfully assembled a total of 106 COI sequences. All assembled COI sequences from this study have been deposited in GenBank with accession numbers OM630610–OM630715.

### COI validation of morphology-based species identification

We analysed assembled COI sequences with BLASTn against the nucleotide collection (nr/nt) database to confirm morphology-based species identification. BLAST analyses revealed 32 cases where top hits indicated a different species identity, taking <95% nucleotide sequence similarity as the threshold to delineate distinct species (Supplementary Table S2). In these cases, the COI sequence of the specimen was then BLAST-aligned against a GenBank record representing the morphological species to verify that the revised identity is a closer match by a significant margin, i.e., more than 2% nucleotide sequence similarity. All species names reported hereafter reflect identities determined by COI barcoding except for cases where COI-based identities were ambiguous, in which case morphology-based identities were retained. In cases where matches were found within a single genus but of multiple species, specimens were indicated as an unknown member of their genus (e.g., *Culex* sp.). Information of the highest-scoring references for all specimens, including details of ambiguous BLASTn results, are recorded in Supplementary Table S2.

Within our COI sequences, we found six unidentified *Culex* species (including two that matched to GenBank entries identified only to the genus level), four unidentified *Mansonia* species, and one unidentified *Mimomyia* species. For *An. baezai*, no existing GenBank records were found at the time this analysis was performed.

### Phylogenetic analysis

Multiple sequence alignment (MSA) were performed on assembled COI and rRNA sequences using the MUSCLE software (Supplementary Files 1–4) (Edgar, 2004; Madeira et al., 2019). As shown in Supplementary Figure S2, the 28S rRNA sequences contain many blocks of highly conserved nucleotides, which makes the result of multiple alignment particularly obvious. We therefore did not test other alignment programs. The multiple alignment of the COI amplicons is even more evident since no gaps are necessary for this alignment.

Phylogenetic tree reconstructions were performed with the MEGA X software using the maximum-likelihood method (S. Kumar et al., 2018). Default parameters were used with bootstrapping with 500 replications to quantify confidence level in branches. For rRNA trees, sequences belonging to an unknown species of parasitic water mite (genus *Horreolanus*) found in our specimens served as an outgroup taxon. In addition, we created and analysed a separate dataset combining our 28S rRNA sequences and full-length 28S rRNA sequences from GenBank totalling 169 sequences from 58 species (12 subgenera). To serve as outgroups for the COI tree, we included sequences obtained from GenBank of three water mite species, *Horreolanus orphanus* (KM101004), *Sperchon fuxiensis* (MH916807), and *Arrenurus* sp. (MN362807).

## DECLARATIONS

### Availability of data and materials

Raw RNA-seq fastq sequence data are available from the corresponding author upon reasonable request. Multiple sequence alignment files are included as supplementary files. All sequences generated in this study have been deposited in GenBank under the accession numbers OM350214– OM350327 for 18S rRNA sequences, OM542339–OM542460 for 28S rRNA sequences, and OM630610–OM630715 for COI sequences (Supplementary Table S1).

## Acknowledgements

We thank members of the Saleh lab for valuable discussions and to Dr Louis Lambrechts for critical reading of the manuscript. We especially thank all medical entomology staff of IP Bangui, IP Cambodge (Sony Yean, Kimly Heng, Kalyan Chhuoy, Sreynik Nhek, Moeun Chhum, Kimhuor Sour and Pierre-Olivier Maquart), IP Madagascar, and IP Guyane for assistance in field missions, laboratory work, and logistics, and Inès Partouche from IP Paris for laboratory assistance. We are also grateful to Dr Catherine Dauga for advice on phylogenetic analyses, and to Amandine Guidez for providing a French Guiana-specific COI reference library.

## SUPPLEMENTARY FILES

**Supplementary Figure S1.**
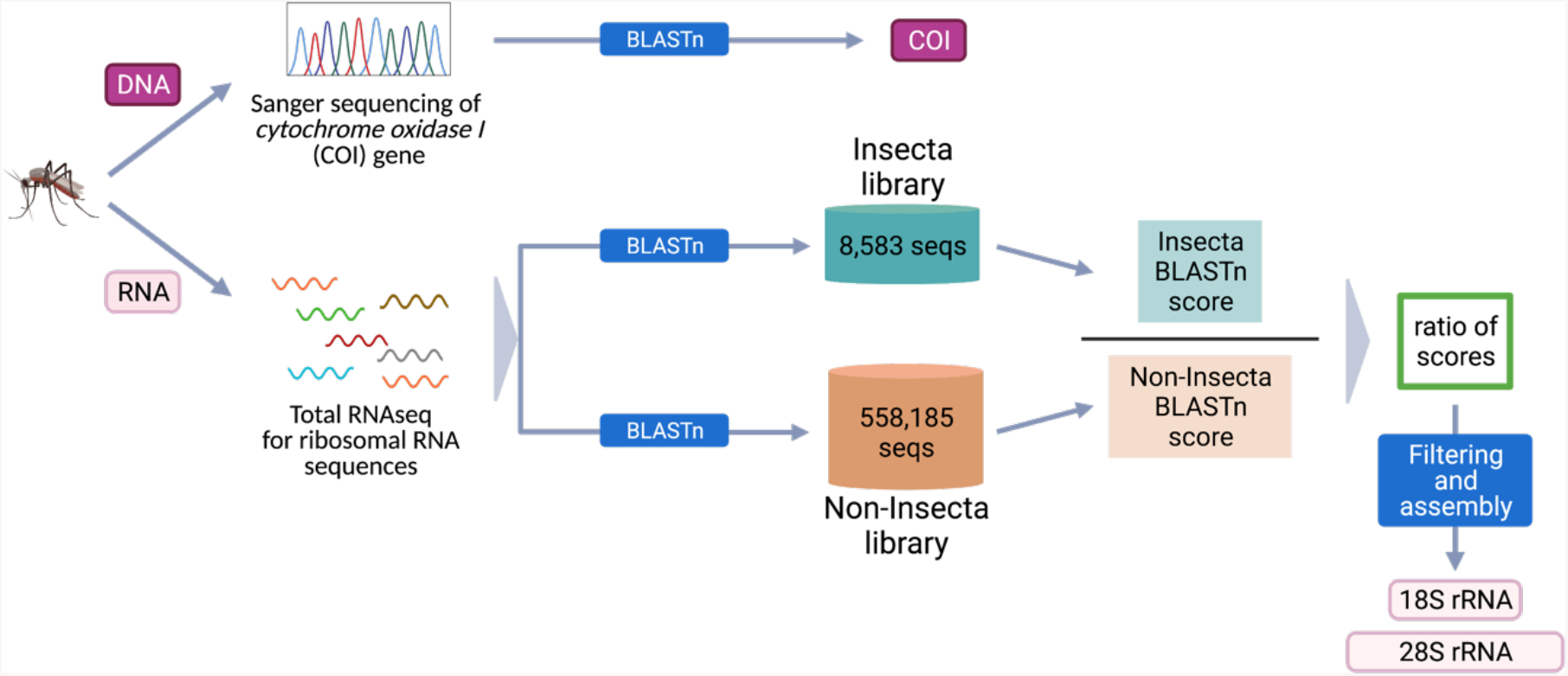
Study workflow from specimens to sequences.

**Supplementary Table S1**. Taxonomic and sampling information on mosquito specimens and associated accession numbers of their 28S, 18S, and COI sequences (XLSX).

**Supplementary Table S2**. COI sequence BLAST analyses summary (XLSX).

**Supplementary Figure S2.**
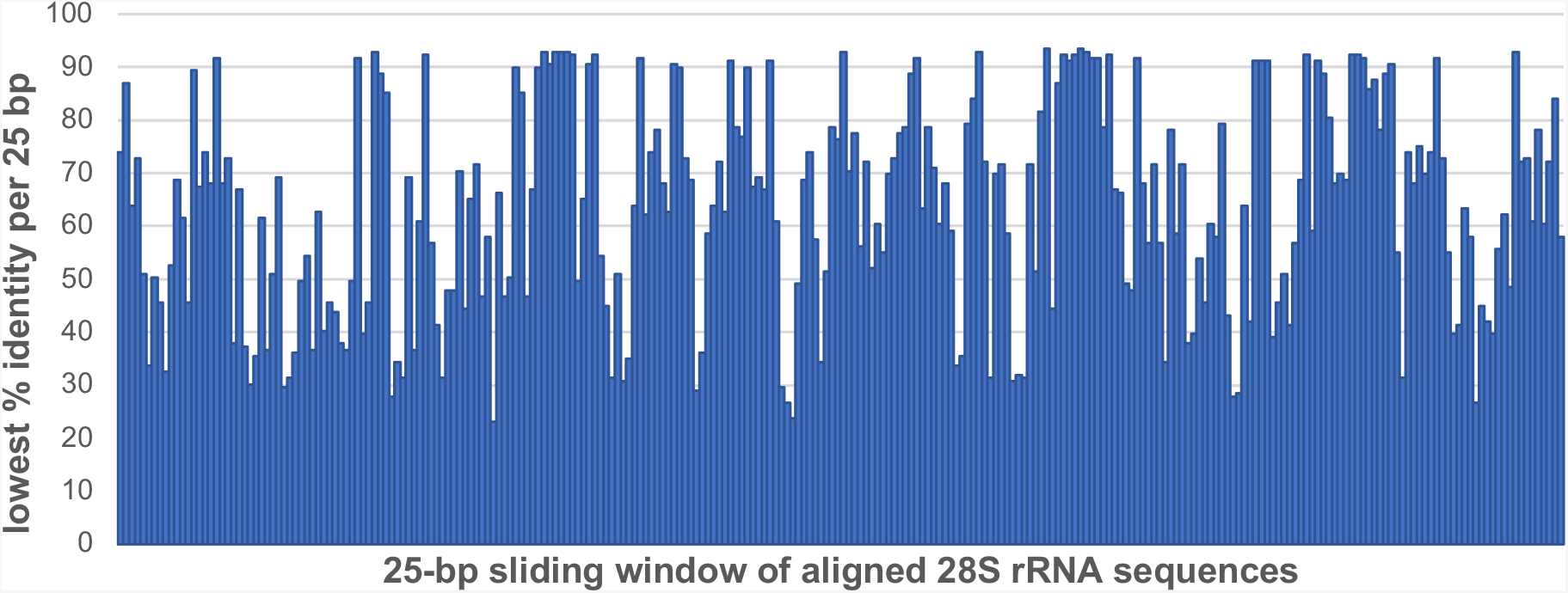
Sequence conservation among 169 28S rRNA sequences obtained from this study and from GenBank combined. Multiple sequence alignment was performed on 28S rRNA sequences, 3900 bp in length. Each bar represents a 25-bp sliding window of the 28S rRNA sequence alignment where the values are the lowest percentage identity found.

**Supplementary Figure S3.**
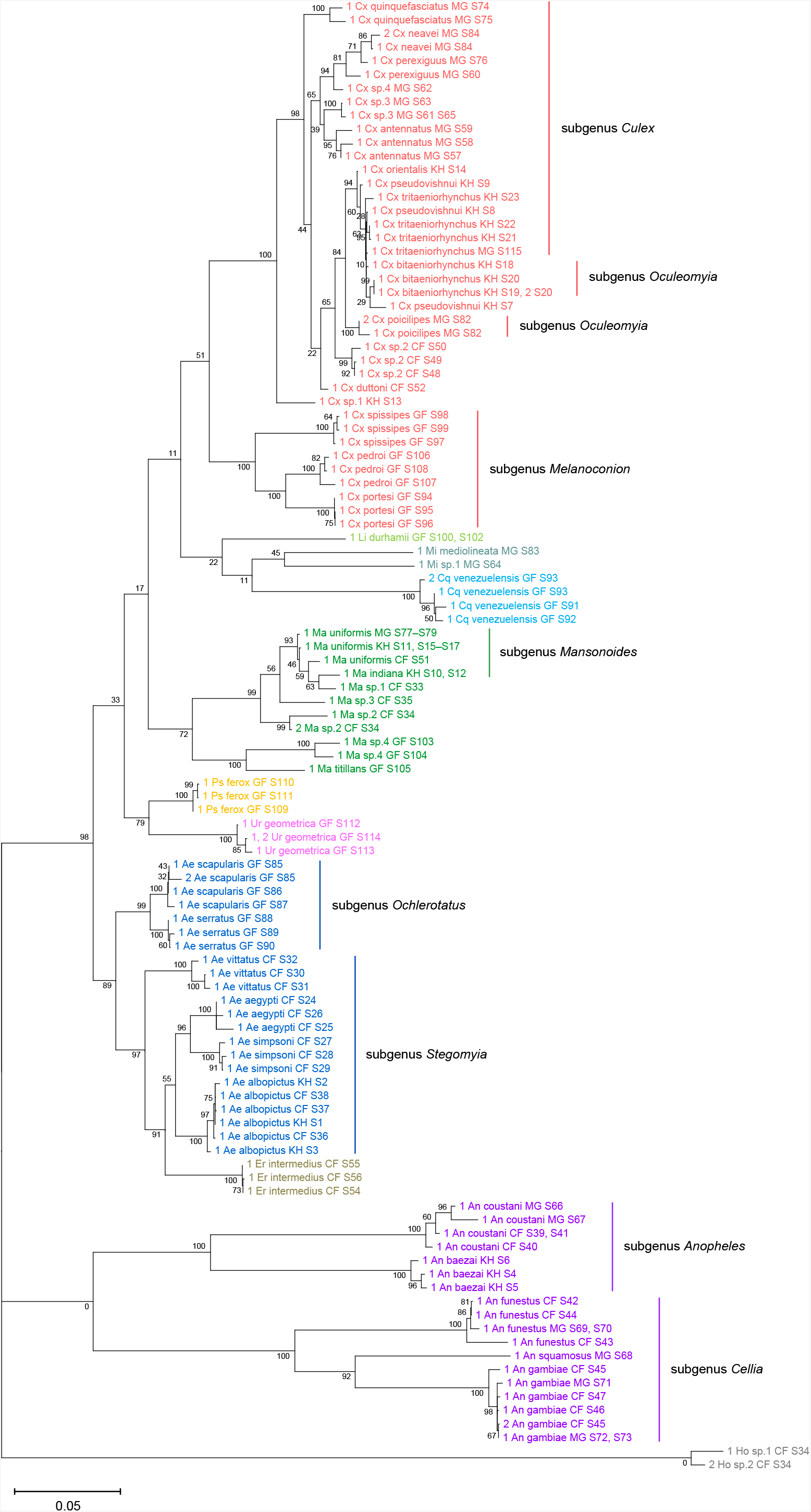
Phylogenetic tree based on 28S sequences generated from this study (3900 bp) as inferred using maximum-likelihood method and constructed to scale in MEGA X (S. Kumar et al., 2018) in (A) and (B) radial format. Values at each node indicate bootstrap support (%) from 500 replication. Each specimen label contains information on taxonomy, origin (as indicated in 2-letter country codes), and specimen ID. Label colours indicate genera: *Culex* in coral, *Anopheles* in purple, *Aedes* in dark blue, *Mansonia* in dark green, *Limatus* in light green, *Coquillettidia* in light blue, *Psorophora* in yellow, *Mimomyia* in teal, *Uranotaenia* in pink and *Eretmapodites* in brown. Scale bar at 0.05 is shown.

**Supplementary Figure S4.**
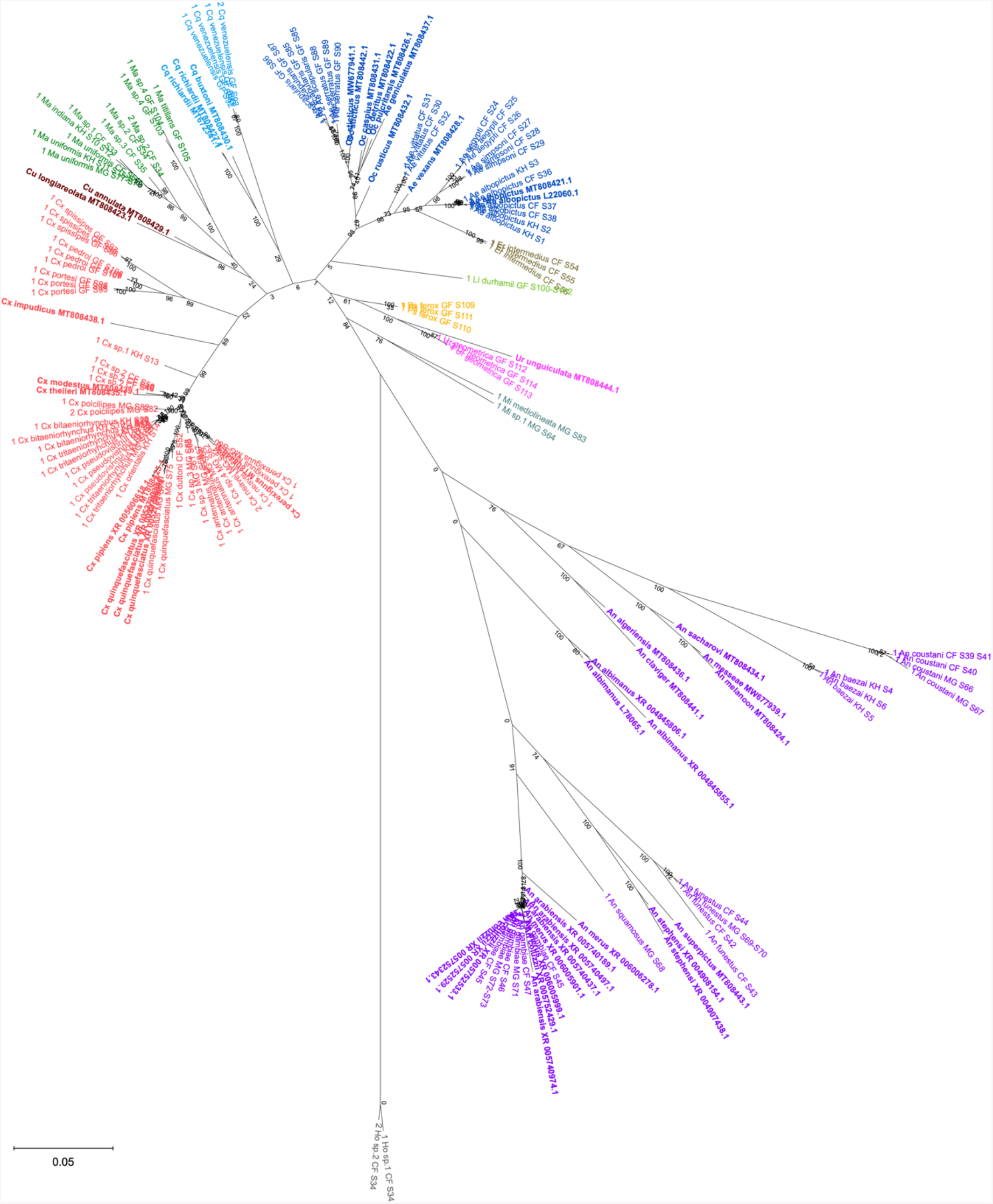
Phylogenetic tree based on 28S sequences generated from this study and from GenBank (3900 bp) as inferred using maximum-likelihood method and constructed to scale in MEGA X (S. Kumar et al., 2018) in radial format to illustrate the distance of Anopheline sequences relative to other Culicidae members. Values at each node indicate bootstrap support (%) from 500 replications. For sequences from this study, each specimen label contains information on taxonomy, origin (in 2-letter country codes), and specimen ID. Labels in bold indicate GenBank sequences with accession numbers shown. Label colours indicate genera: *Culex* in coral, *Anopheles* in purple, *Aedes* in dark blue, *Mansonia* in dark green, *Culiseta* in maroon, *Limatus* in light green, *Coquillettidia* in light blue, *Psorophora* in yellow, *Mimomyia* in teal, *Uranotaenia* in pink and *Eretmapodites* in brown. Scale bar at 0.05 is shown.

**Supplementary File 1:** Multiple sequence alignment of 169 28S rRNA sequences from this study and from GenBank (FASTA).

**Supplementary File 2:** Multiple sequence alignment of 122 28S rRNA sequences, including two sequences from *Horreolanus sp*. (FASTA).

**Supplementary File 3:** Multiple sequence alignment of 114 18S rRNA sequences, including two sequences from *Horreolanus sp*. (FASTA).

**Supplementary File 4:** Multiple sequence alignment of 106 COI sequences (FASTA).

