## Supplementary figures and images for "Decoding rRNA sequences for improved metagenomics of sylvatic mosquito species"

### Supplementary Figure S1

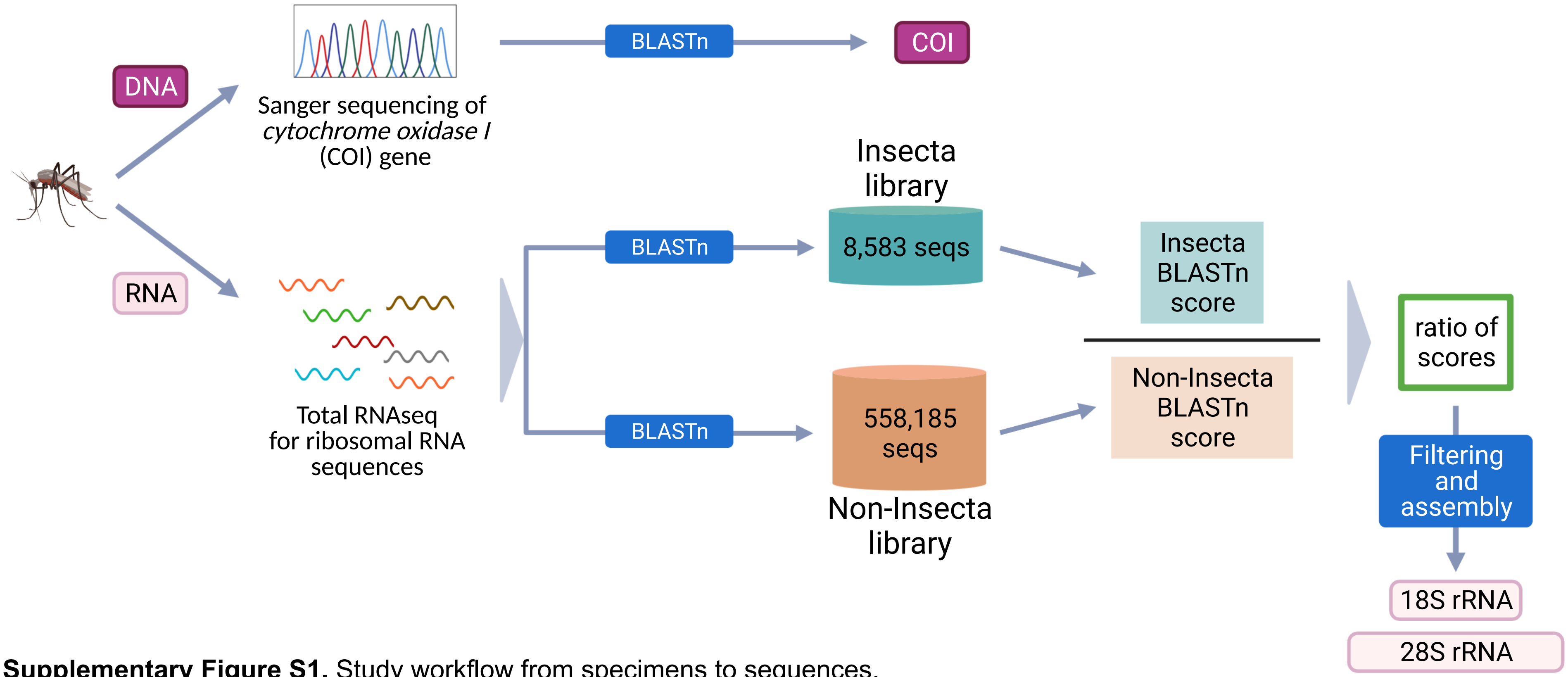


**Supplementary Figure S1.** Study workflow from specimens to sequences.

### Supplementary Figure S3

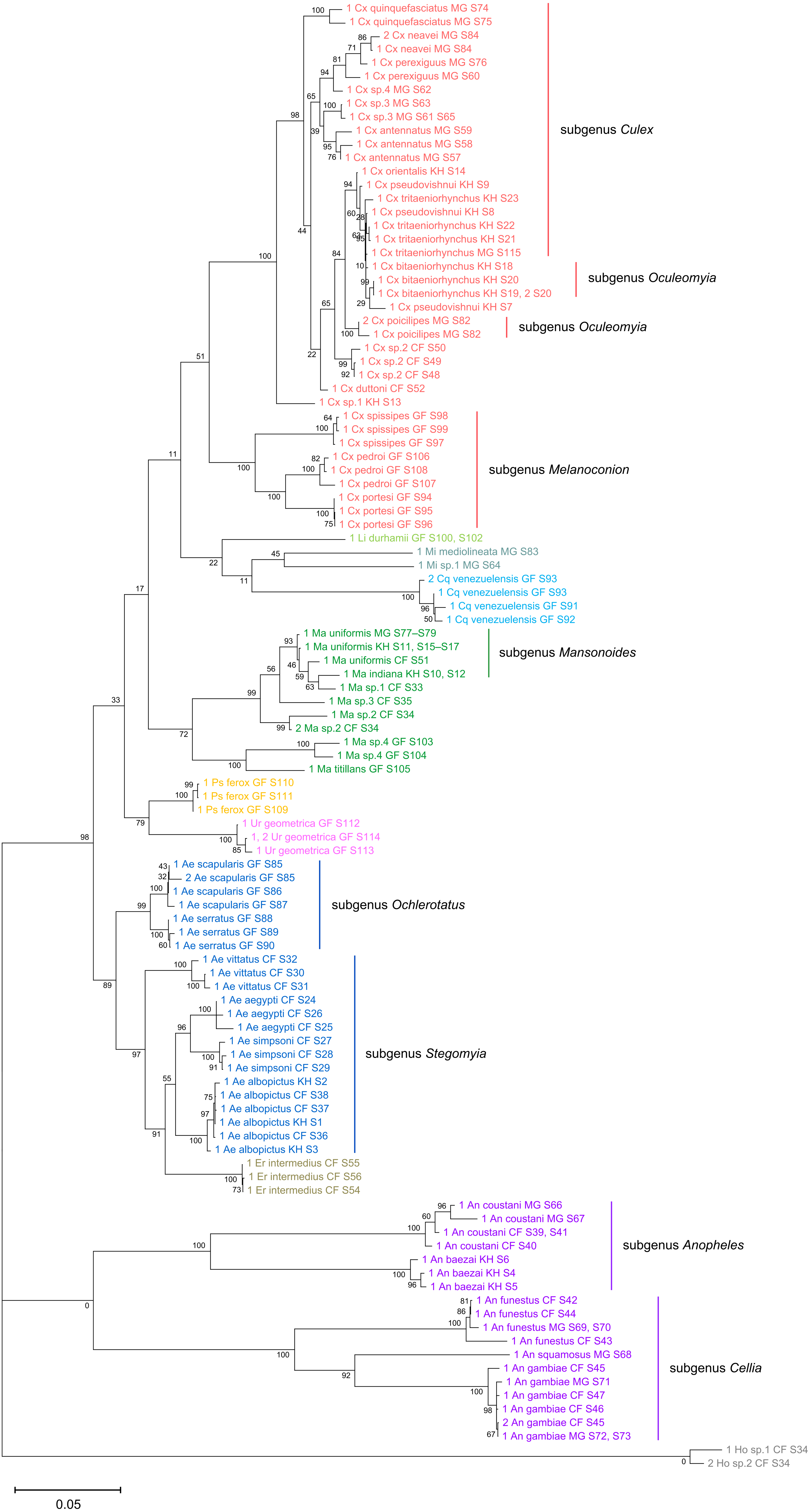

### Supplementary Figure S4

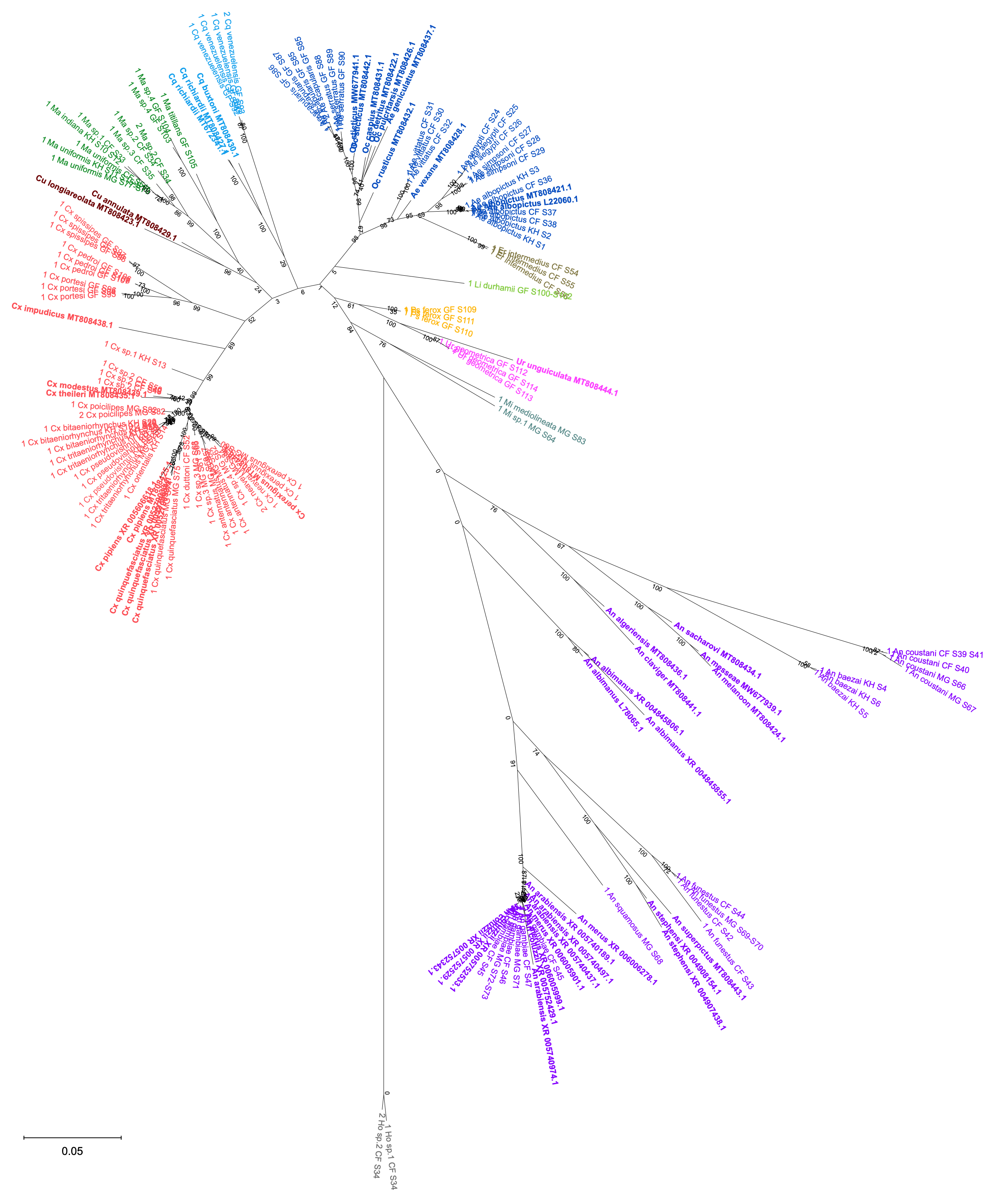
