## Supplementary Figure S2 for "Decoding rRNA sequences for improved metagenomics of sylvatic mosquito species"

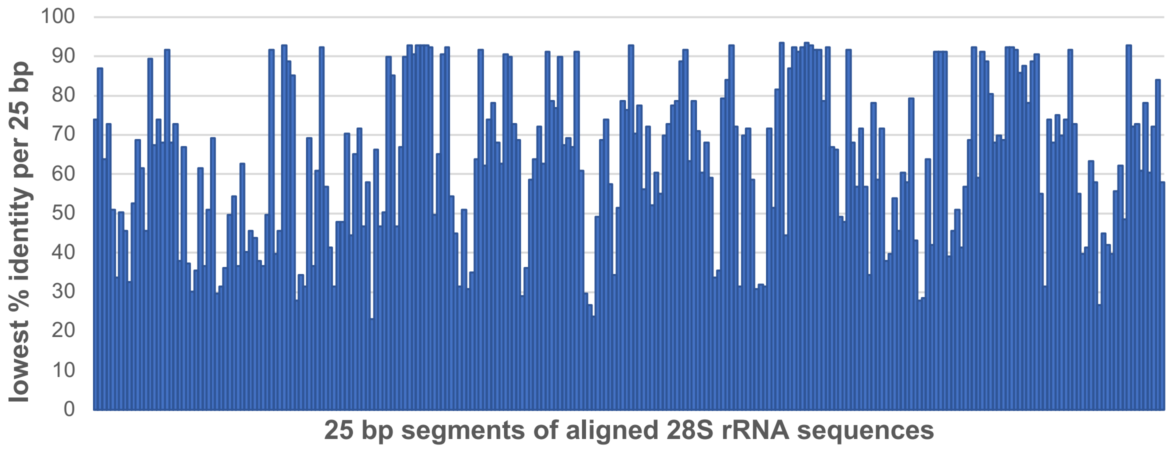


**Supplementary Figure S2.** Sequence conservation among 169 28S rRNA sequences obtained from this study and from GenBank combined. Multiple sequence alignment was performed on 28S rRNA sequences, 3900 bp in length. Each bar represents a sliding 25-bp segment of the 28S rRNA sequence alignment where the values are the lowest percentage identity found.

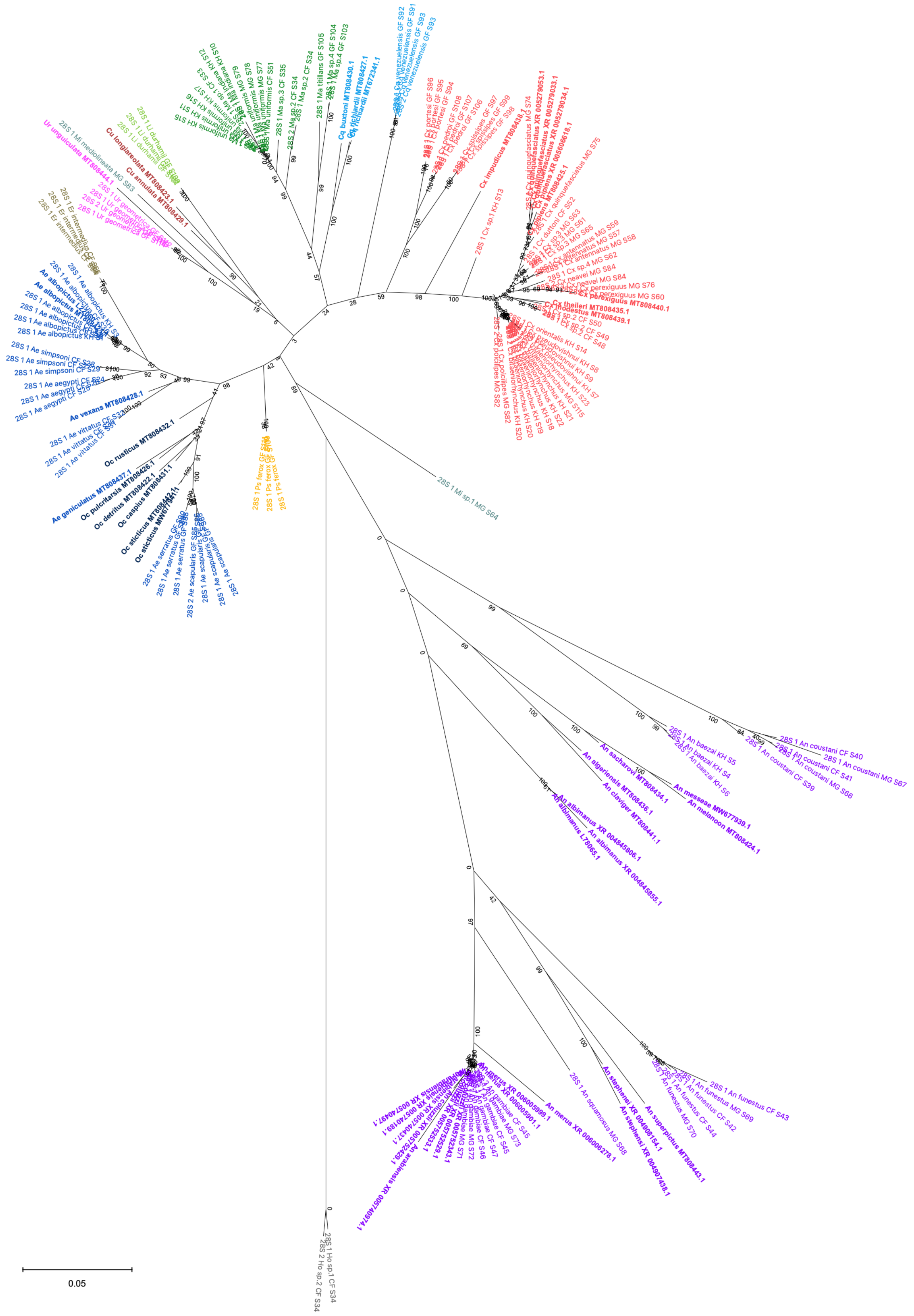

**Supplementary Figure 2.** Phylogenetic tree based on 28S sequences generated from this study and from NCBI databases combined (3900 bp) as inferred using maximum-likelihood method and constructed to scale in MEGA X in radial format (18). Values at each node indicate bootstrap support from 500 replications. For sequences from this study, each specimen label contains information on its taxonomy, origin (as indicated in 2-letter country codes), and specimen ID. Labels in bold indicate sequences derived from NCBI. Label colours indicate genera: *Culex* in coral, *Anopheles* in purple, *Aedes* in dark blue, *Mansonia* in dark green, *Culiseta* in maroon, *Limatus* in light green, *Coquillettidia* in light blue, *Psorophora* in yellow, *Mimomyia* in teal, *Uranotaenia* in pink and *Eretmapodites* in brown. Scale bar at 0.05 is shown.
